# AI in paleontology

**DOI:** 10.1101/2023.08.07.552217

**Authors:** Congyu Yu, Fangbo Qin, Akinobu Watanabe, Weiqi Yao, Ying Li, Zichuan Qin, Yuming Liu, Haibing Wang, Qigao Jiangzuo, Allison Y. Hsiang, Chao Ma, Emily Rayfield, Michael J. Benton, Xing Xu

## Abstract

Accumulating data have led to the emergence of data-driven paleontological studies, which reveal an unprecedented picture of evolutionary history. However, the fast-growing quantity and complication of data modalities make data processing laborious and inconsistent, while also lacking clear benchmarks to evaluate data collection and generation, and the performances of different methods on similar tasks. Recently, Artificial Intelligence (AI) is widely practiced across scientific disciplines, but has not become mainstream in paleontology where manual workflows are still typical. In this study, we review more than 70 paleontological AI studies since the 1980s, covering major tasks including micro-and macrofossil classification, image segmentation, and prediction. These studies feature a wide range of techniques such as Knowledge Based Systems (KBS), neural networks, transfer learning, and many other machine learning methods to automate a variety of paleontological research workflows. Here, we discuss their methods, datasets, and performance and compare them with more conventional AI studies. We attribute the recent increase in paleontological AI studies to the lowering bar in training and deployment of AI models rather than real progress. We also present recently developed AI implementations such as diffusion model content generation and Large Language Models (LLMs) to speculate how these approaches may interface with paleontological research. Even though AI has not yet flourished in paleontological research, successful implementation of AI is growing and show promise for transformative effect on the workflow in paleontological research in the years to come.

**Highlights:**

- **First systematic review of AI applications in paleontology.**
- **There is a 10 to 20-year gap between AI in paleontology and mainstream studies.**
- **Recent progress in paleontological AI studies is likely a result of lowering bar in training and deployment.**
- **Future direction discussed for interactions between paleontology and AI.**

## 1. Introduction

### 1.1 Data-driven earth sciences

Recently, artificial intelligence (AI) has shown fast-growing applications in a wide range of fields in earth sciences and demonstrates the transformation towards data-driven studies (Bergen et al. 2019; Reichstein et al. 2019; Cheng et al. 2020; Sun et al. 2022). Global hydrology (Yao et al. 2023), weather forecasting (Bi et al. 2023; Zhang YC. et al. 2023), seismology (Kong Q. et al. 2019, Mousavi & Beroza 2023), carbon cycle (Tao et al. 2023), and many other subfields in earth sciences have benefited from advances in AI. However, due to challenges in data collection and processing, task complexity, lack of proper models, a large number of earth science fields, including paleontology, have primarily relied on less automated, more manual workflows.

Studies of deep-time biodiversity are mostly based on incomplete fossil records. Large-scale datasets and complex statistical methods have enabled deep-time high-resolution evolutionary models of both global and local biodiversity from marine and terrestrial faunas (Fan J. et al. 2020; Zhou et al. 2021; Dai et al. 2023), and also offer an indispensable opportunity to evaluate the causes and processes of a series of bio-geological events. Reconstructing the co-evolution of life and environment through geological time is a key to understand earth history. Specifically, the "Big Five" mass extinction events have been observed across the Ordovician–Silurian, late Devonian, Permian–Triassic, Triassic–Jurassic, and Cretaceous–Paleogene boundaries (Sepkoski 1978, 1979, 1984; Payne and Finnegan 2007), with profound perturbations in biogeochemical processes. While previous studies have taken a diversity of geochemical and modeling approaches to reveal the evolution across time and space, the ability to integrate various approaches to provide comprehensive analysis and underlying mechanism is still limited. Recently, the paleontological community started to pay attention to data-driven studies that combine large-scale datasets with AI tools. For sedimentary studies, Emmings et al. (2022) applied text mining coupled to multivariate statistical analysis over a large library of published sedimentary datasets from the Precambrian to present to expand the application of pyrite morphology and chemistry as paleo-redox proxies. For ocean chemistry studies, Mete et al. (2023) developed Gaussian Progress Regression machine learning models to accurately simulate the spatially and vertically distribution of trace elements (e.g., barium) in the sea.

However, due to a scarcity of samples from certain temporal of spatial regions, enormous gaps exist and are often approximated by extrapolating disparate datasets, which might average out finer changes that might be present. To date, systematic statistical analysis has not yet been completed based on big data and there are few common datasets for model evaluation? Therefore, it is difficult for machine to learn from current earth science data. Comparing with other earth science fields, there are far fewer AI-based paleontological studies, and even fewer for vertebrate paleontology. In this study, we show the development of AI-based paleontological studies since the 1980s, covering distinctive tasks and groups of organisms, with a particular focus on the evolution of datasets and algorithms.

### 1.2 Data-driven paleontology

Since paleontology studies ancient organisms, it is primarily based on fossil materials. However, fossil records are extremely patchy fragments from evolutionary history, and large-scale quantitative paleontological studies have only been feasible when relevant datasets are compiled. A large portion of traditional paleontological studies focus on fossil morphology, including anatomical description and comparative studies among specimens, which can be considered as “fossil-driven” studies. On the other hand, “data-driven” paleontological studies can be characterized as studies that work with relatively large number of fossil specimens, and apply various analytical techniques to discover patterns from data. Ultimately, all paleontological studies are based on fossil specimens, thus the boundary between “fossil-driven” and “data-driven” cannot always be well defined. Consequently, here we make no attempt to accurately define what “fossil-driven” and “data-driven” paleontological studies are. However, we observe that progress in paleontological data collection, application of advanced analytical methods, and increasing availability of data sharing and distribution, have resulted in an increase in the number of paleontological studies that incline to the “data-driven” paradigm.

Early example of “data-driven” paleontological studies can be traced back to Matthew (1926), who studied horse macroevolution by compiling the fossil records of horses and their early relatives, globally and systematically comparing morphological changes in teeth, limbs and skulls, temporal and spatial distribution, and tempo and mode of their evolution. Although quantitative analysis was lacking in Matthew (1926), such systematic summary and comparison of horse evolution presented a primitive form of data-driven paleontological research. More systematic work with a much broader scope about evolution from the Precambrian to the Phanerozoic was summarized by Simpson (1944). Simpson (1944) was one of the first to present a comprehensive plan to convert paleontological data into numerical, statistical problems to explore rates and modes of evolution, giving numerous examples from his favorite group, the horses, exploring evolution of single and multiple morphological traits.

Later, with much more abundant fossil records, historical marine biodiversity became a hotspot in data-driven paleontological studies. A classic example is the series of works on Phanerozoic marine biodiversity and major extinction events by Sepkoski (1978, 1979, and 1984) and Raup & Sepkoski (1982). On the other hand, quantitative paleontology has been fast developing since the middle 20^th^ century. Numerical taxonomy (*sensu* phenetics) was proposed in late 1960s to build a workflow that is capable of quantitatively comparing characteristics across taxa to reconstruct their evolutionary history (Sneath & Sokal 1962, 1973). Numerical taxonomy introduced a series of quantitative methods to taxonomic studies that tried to eliminate influences from intuitive judgements and allow studies to be reproducible. Along the same line, the study of cladistics developed by Hennig (1966, 1999) at almost the same time. While both phenetics and cladistics use morphological characters as data, the former clusters taxa based on overall morphological similarity while the latter clusters taxa based on synapomorphies, or shared derived character states. The use of phenetics has been superseded by cladistics, which is still used in paleontological studies to infer phylogenies along with other statistical phylogenetic frameworks (i.e., maximum likelihood and Bayesian methods).

Paleontology can hardly work with molecular data that provide ample information for life science studies, and the yet oldest ancient DNA sample sequenced yet was only traced back to 2.4 MYA (Kjær et al. 2022), which is incredibly young by geological standards and as such aDNA only covers a tiny sliver of evolutionary history. Although attempt have been made to establish the so-called “paleo-bioinformatics”, which is derived from “bioinformatics”, on the basis of information theory and morphological data (Yu et al. 2021). In general, fossil morphology is the core resource for a large portion of paleontological studies. The compilation of morphological data has made more quantitative studies feasible and the introduction of morphospace and other quantitative tools. Raup (1966) analysed the shape of different shell forms by illustrating specimens in a space given by three geometric measurements and now is often seen as the primitive form of morphospace (Budd 2021). During the decades since these works, methodological advancements in quantitative characterization, visualization, and analysis of biological form has truly revolutionized phenotypic research, including paleontology. Chief among these advances is the Procrustean paradigm in geometric morphometrics (GM), which uses Cartesian coordinates to mathematically describe differences and changes in shape (Bookstein 1992; Rohlf & Marcus 1993; Slice 2007; Adams et al. 2013). Coupled with non-destructive 3D imaging techniques in fossils, GM is common practice in quantitative comparative morphological studies and has been widely used in various groups of organisms such as dinosaurs (Bhullar et al. 2012; Hanson et al. 2021; Choiniere et al. 2021) and synapsids, including mammals (Lungmus & Angielczyk 2019; Goswami et al. 2022). These highly multivariate, or multidimensional, anatomical data have allowed paleontologists to investigate the tempo and mode of phenotypic evolution with much greater morphological fidelity.

Due to the scope of this study, we cannot list every kind of quantitative paleontological study here, there are also many more textbooks and software available (e.g., Hammer & Harper 2001, 2008). Even though the amount, modalities, and scopes of available fossil data have rapidly increased during the last six decades, and we have witnessed a variety of data-driven paleontological studies on diversified topics and organisms, paleontological AI applications to date are scarce. The majority of paleontological studies still rely on largely manual workflows for fossil preparation, morphological description, and downstream analyses.

### 1.3 A brief history of AI and classic tasks

AI comprises a broad range of techniques that enable a computer to process data, execute tasks, and even learn from experience, which simulates human intelligence. Thus, AI systems are expected to assist or replace humans in various intellectual tasks with high performance. The major advantage of an AI system compared to a human is the high throughput and ability to perform consistent and accurate data processing. Typical applications of AI include face recognition, object detection, natural language processing, machine translation, speech recognition, content generation, video game development (e.g., procedural-content generation), medical diagnosis, etc.

Although ideas of machine-aided analyses had been suggested earlier, AI was first proposed as a scientific idea at a workshop held in Dartmouth College (New Hampshire, USA) in 1956, which was the groundbreaking landmark of this field (McCarthy et al. 2006). There have been ups and downs in the development of AI since 1956 following advances in hardware and algorithms and mismatches between expectation and reality. During its growth, novel ideas in AI have been frequently proposed and rejected. For example, the Knowledge Based System (KBS) was once popular but has now mostly been abandoned (Bell 1985). There have been several reviews of the history of AI and its subfields such as deep learning (LeCun et al. 2015; Jordan & Mitchell 2015; Baranjuk et al. 2020; Bengio et al. 2021; Toosi et al. 2021; Xu YJ. et al. 2021).

Since traditional, non-AI models have very limited adaptability, complexity, and optimality, machine learning is the current mainstream AI theory, which allows a user to optimize its performance on a certain task by learning from experience through provision of training data sets. The learning paradigms include supervised learning, semi-supervised learning, unsupervised learning, and reinforcement learning (Fig. 1). Typically, a machine learning model (e.g., Gaussian mixture model, support vector machine (SVM), neural network, random forest, etc.), is trained on an input dataset, during which the model’s output is evaluated at each iteration according to ground truths and the model’s parameters are gradually tuned according to the learning objective. The data formats of the input and output can be in various modalities, while the common problem is to predict discrete or continuous values that represent some certain information. Therefore, the inputs and outputs can be formalized as map, tensor, vector, probabilistic distribution, etc., so that the informative values can be calculated and predicted.

**Figure 1.**
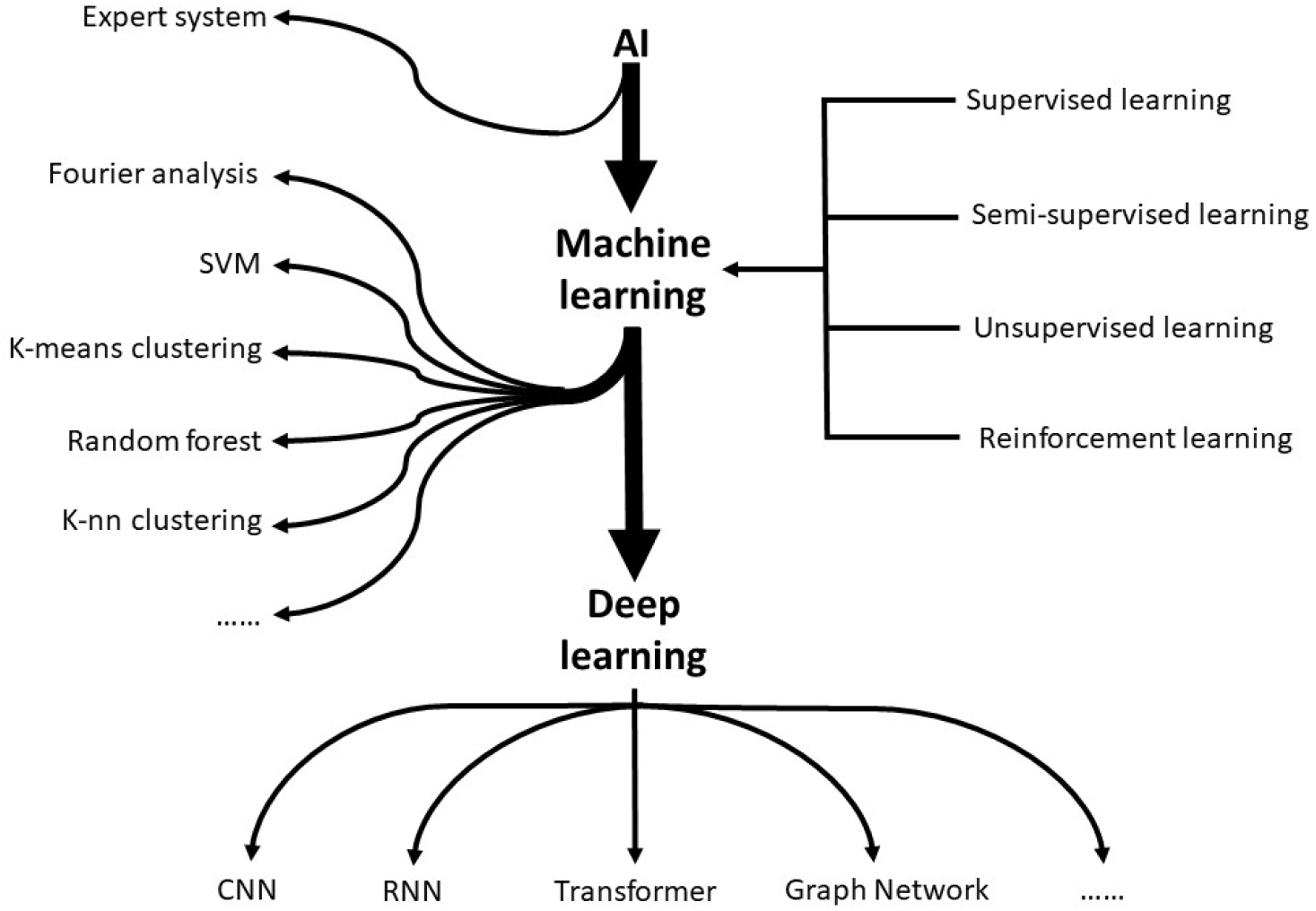
A simplified structure of AI.

Deep learning (DL) is a subfield of machine learning methods that started with Hinton et al. (2006), Hinton & Salakhutdinov (2006) and Bengio & LeCun (2007), but its core idea was presented much earlier (e.g., LeCun 1989). With the availability of large training datasets and hardware, DL has boomed since 2012 (Krizhevsky et al. 2012; Goodfellow et al. 2014). DL is exemplified by deep neural networks (DNN) with multiple layers (up to >100) and representation learning, which allows a DNN to learn to perceive rich, complex, and hierarchical feature representations from unstructured data. For example, given an image, a DNN can extract the low-level features (color, texture, edge, etc.), the mid-level features (shape, parts, etc.) and high-level features (semantic, category, context, etc.). Compared to traditional ML techniques, DL has absolute superiority in description and generalization abilities, especially when large datasets with annotation are available for training.

## 2. Paleontological AI study development

In this section, we review paleontological AI applications from the early 1980s to 2023. Studies are categorized by their tasks, including classification (micro and macro fossils), segmentation, and prediction (Fig. 2A). Although the organisms covered here are from distantly related groups, the central idea and methods used in research are usually interchangeable. For each section, we make efforts to arrange reviewed studies chronologically to show the rise and fall of different AI methods and advancements in fossil data compilation. We further divide section 2.1 on microfossil classification into four sub-sections since microfossils have dominated past paleontological AI studies and they represent the most complete history of AI applications in paleontology.

**Figure 2.**
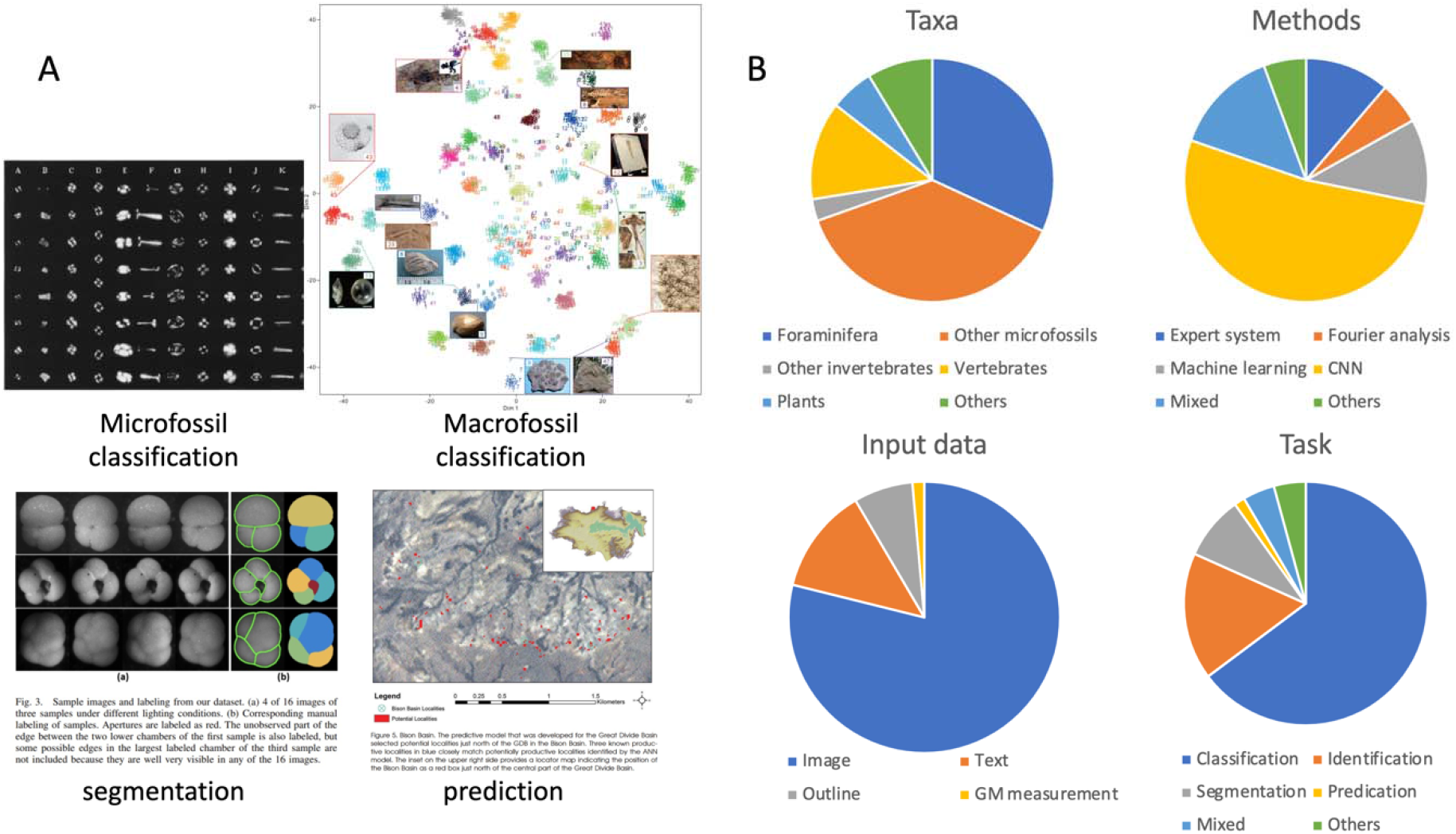
A. Examples of paleontological AI tasks, images are modified from Doffus & Beaufort (1999) Microfossil classification, Liu X. et al. (2023) Macrofossil classification, Ge et al. (2017) segmentation, and Anemone et al. (2011) prediction. B. proportions of taxa, methods, input data, and task in paleontological AI studies.

### 2.1 Microfossil Classification

Microfossils have always been a subject of intensive paleontological study due to their large quantity (resulting from high preservation potential) and imperative roles in evolution, sedimentary geology, paleoclimate, and many other fields. In contrast with macrofossils, microfossils generally have a size less than 1 mm that can be barely seen by the naked eye. Microfossils collectively refer to a variety of organisms including but not limited to foraminifera, conodonts, algae such as coccoliths, plant pollens, and fossil fragments from other organisms (e.g., ichthyoliths). The recognition, identification, and classification of microfossils is normally tedious, and thus many researchers have proposed methods to automate different aspects in a traditional handcrafted research workflow, such as sampling, imaging, measurement, identification, classification, etc. The most fundamental challenge is microfossil classification, including the classification from similarly sized particles and into different species. Many early attempts applied *Fourier analysis* to extract patterns from outline shapes and other components of shape that are diagnostic for species identification (Healy-Williams 1983, 1984) or to identify fossils (Belyea and Thunell 1984; Burke et al. 1987; Garratt 1992; Garratt and Swan 1992). Thresholding based on image pixel gray values was also used in outline extraction (segmentation) from fossil images (Hills 1988). However, these early methods can hardly deal with realistic scenarios, as they often require extensive human effort in data preparation or can only be applied to extremely specific tasks.

#### 2.1.1 Knowledge Based System for Microfossil Classification

As an early stage in AI, automated specimen identification or classification in paleontology can be traced back to studies that constructed microfossil datasets at various scales and tried to infer the taxonomic status of either recorded or new specimens (Brough & Alexander 1986, Beightol & Conrad 1988, Riedel 1989, Swaby 1990 & 1992, Athersuch et al. 1994, Liu S. et al. 1994) and Yu S. et al. 1996). They used a branch of AI called *Knowledge Based System* (KBS, or *Expert System*) that encompasses a knowledge base and an inference engine. In fossil KBS, there was usually a database of annotated subjects, such as fossil specimen images, and a set of pre-determined discriminate rules. In such a system, new specimens can be identified or classified based on given rules (e.g., the presence or absence of a particular feature, the position of a certain anatomical structure) and the rules themselves may be modified and updated. Most of these KBS are based on text rules and may store a certain number of images for illustration reference. Sometimes the size of a KBS can be fairly large, such as Swaby (1990, 1992) who built visual identification expert systems (VIDE) for fossil foraminifera and conodont identification, which integrated 3,500 images, 100 attribute-value tables, and over 10,000 lines of text information. Liu S. et al. (1994) constructed a dual-step identification system that embedded a KBS and an image analysis subsystem, which extracted graphic information from foraminifera chambers and suture images in addition to previous shape analysis based on Fourier analysis and edge detection. The system was built in CLASSIC, a knowledge-based system shell that provides the knowledge base scheme and inference engine. Most KBS were built up according to a given taxonomic hierarchy and anatomical terminologies, resembling a traditional paleontological key, such as a dichotomous key, in using diagnostic characters for classification. However, the scope of those studies was usually incomplete for more general identification; for example, there were only 30 identifiable species of modern planktonic foraminifera (out of a total of ∼50) with 25 discriminate rules in another CLASSIC based KBS prototype by Yu S. et al. (1996). Further, the limited computing performance in the 1990s meant that KBS often required considerable time in identification (e.g., 3∼4 min/specimen by Yu S. et al. 1996).

Most KBS studies in paleontology were conducted from the late 1980s to mid-1990s, and a few studies continued to explore such application in the 2000s. For example, Kaya et al. (2013) built an expert system to classify pollen images of *Onopordum*, an extant plant in the family Asteraceae. The community has witnessed a similar rise and fall of KBS in conventional AI studies (Davis 1982; Bell 1985). As a pre-designed system, KBS is commonly restricted by the prior knowledge from experts that almost inevitably incorporates personal biases and, although robust in processing recorded inputs, KBS cannot process unfamiliar objects. It thus has limited intelligence (though its definition is ambiguous) or robustness. The capability of a given system may be improved by incrementally adding “knowledge” and discriminate rules, but ultimately KBS has to be built on well-organized databases and a confidently agreed taxonomy, both of which are not practical for extinct and extant organisms. KBS cannot “learn” or “discover” hidden knowledge, thus its popularity has gradually declined as other probability-based methods (e.g., random forest) perform better in prediction and require less effort in design and maintenance. However, the modern knowledge-graph may be seen as a successor to KBS by incorporating knowledge from a given field or even a very broad range (e.g., Wikipedia), although the reasoning part remains neglected to a great extent.

#### 2.1.2 (Convolutional) Neural Network for Microfossil Classification

The concept of a neural network is now a core method in AI studies. It arranges interconnected nodes in a layered structure that resembles neurons in the brain and can approximate any given function at reasonable cost (Hornik 1989). Convolutional Neural Networks (CNN) are the most commonly used neural network in the computer vision area, which uses the convolution operation to extract features from the input and has shown outstanding performance in image-related tasks (Krizhevsky et al. 2012). The initial attempt to deploy CNN in image processing was conducted by Fukushima (1980), and its paleontological use can be traced back to *Système de Reconnaissance Automatique de Coccolithes* (SYRACO, or SYRACO1) by Dollfus & Beaufort (1996), close on the heels of the application of neural networks to identification of extant marine phytoplankton (Boddy et al. 1994, firstly used back-propagation to update neuron status in the network, a commonly used training method). Coincidently, this was when one of the last paleontological KBS studies was conducted (Yu S. et al. 1996). SYRACO used two-dimensional fast Fourier transformation and a two-layered neural network to identify planktonic foraminifera images; a detailed description of the system is given by Dollfus (1997). Its updated version SYRACO2 was introduced by Dollfus & Beaufort (1999), which used a deeper 5-layered neural network and back-propagation algorithm. On the basis of larger training datasets including 2,000 images from 13 coccolith species and non-coccolith objects, SYRACO2 showed overwhelmingly better performance than its predecessor. SYRACO2 can make 40 classifications per second on the basis of a 200 MHz processor and has reached a mean recognition efficiency of 86%, while SYRACO1 was 49%. Due to the significantly increased number of parameters (∼800,000) in SYRACO2 at that time, Dollfus & Beaufort (1999) also named it a fat neural network. Beaufort & Dollfus (2004) added parallel neural networks and dynamic view to improve the performance in object detection and classification accuracy. The SYRACO dynamic version added five motor modules for data argumentation by correcting translation, rotation, dilatation, contrast, and symmetry in fossil images. However, as the analysis was run on sedimentary samples that contain numerous coccolith-sized objects, there was a real problem for SYRACO through its iterations of false positives from the inclusion of similar objects. The dynamic version was probably not an ideal update as CNN itself is space invariant, therefore four of the five modules were theoretically not necessary. Nevertheless, the appearance and continuous iteration of SYRACO indicate that the deficiencies of KBS in automated paleontological workflow had been realized, while the application of CNN continued to unfold in both paleontological studies and other fields. The SYRACO system has been actively used in analyzing fossil coccoliths from marine sediments since its first deployment (Beaufort et al. 2001, 2022), which has been one of the most successful and long-lasting applications among paleontological AI studies.

Schiebel et al. (2003) and Bollmann et al. (2004) gave a brief review on automated sedimentary particle (including foraminifera) analysis although there were only a few automated micropaleontological studies at that time. They explained the need to build microscope systems to automatically acquire fossil images in the field, and pointed out the differences between structural/statistical techniques and Artificial Neural Network (ANN) in automated image analysis. The first category requires given knowledge from experts (e.g., KBS), but the tailored features and inherent inflexibility make those techniques incapable of working with new taxa or messy sediments. At that time, ANN was more likely to be a promising direction according to the results from pollen (France et al. 2004) and coccoliths (Dollfus & Beaufort 1996, 1999; Beaufort & Dollfus 2004). Schiebel et al., (2003) and Bollmann et al. (2004) developed the Computer Guided Nannofossil Identification System (COGNIS) for the segmentation, preprocessing, and classification of microfossil images. The training dataset included 979 images covering 14 Holocene coccolith species at a resolution of 48 × 48 pixels and a classification dataset of 715 images from the same 14 species. Their 5-layered CNN reached a recognition rate of 75% on average, but the error rates varied significantly across species from 3% to 88%. The low resolution in this study seems to have been a compromise between computational cost and sampling, and the 48 × 48 pixels resolution came from down-sampling of the original image. COGNIS-light was developed at the same time to classify only a particular foraminifera species *Florisphaera profunda* from all other biotic or abiotic particles. The training dataset had 1000 images of *F. profunda* and 1000 images of other particles, but such a training strategy resulted in strong false positives, likely due to the strongly biased samples.

Tetard et al. (2020) used ResNet-50 as well as other models to identify fossil radiolarians. Based on a training dataset with 21746 images classified in 132 classes, their model reached an overall accuracy of about 90%. However, the automated workflow from image acquisition to segmentation and recognition could take as long as an hour per sample, and more vitally, its application to images from other datasets without preprocessing led to significant lower accuracy, indicating that the model was significantly overfit to the training dataset. Itaki et al. (2020a) constructed the microfossil Classification and Rapid Accumulation Device (miCRAD) for automated single species recognition based on CNN, which was capable of classifying two radiolarian species from others with an accuracy exceeding 90%. Further, the authors suggested that such a model could be used to classify other microfossils such as foraminifers and diatoms if properly fine-tuned. Itaki et al. (2020b) expanded the dataset and focused on a widely distributed Pleistocene species, *Cycladophora davisiana,* and the results had high consistency with a human expert while saving significant time.

In summary, from the late 1990s to early 2000s, CNN made significant progress in paleontological AI studies and resulted in two successful applications, SYRACO and COGNIS. CNN based models require much less effort in system construction and also run much faster than KBS but do require significant front-loaded time input to generate high-quality training datasets.

#### 2.1.3 Other Machine Learning Methods for Microfossil Classification

While CNN-based models began to thrive in paleontological AI studies, other machine learning methods still played essential roles, especially when the data scale was small. Many studies combined traditional machine learning methods and neural network in fossil related tasks. Marmo and Amodio (2006) and Marmo et al. (2006) used k-Nearest Neighbor (k-NN) and a three-layered perceptron classifier to automatically classify chamber arrangements in foraminifera. However, both the training (207 and 200 images) and testing datasets (70 and 80 images) comprised too few images and only five classes, and the extremely high accuracy (97.1%) from the perceptron classifier is likely a result of over-fitting. Wong (2011) developed Microfossil Quest, an interactive system for microfossil search, identification, and education. This system integrated KBS and various machine learning methods including k-NN, K-means clustering, ANN, etc. Almost 4,000 specimens were included in the database and many of the identifications were collected using crowdsourcing; however, the testing experiment only used 238 specimens and achieved accuracy at the genus and species level of 81% and 47%, respectively. The concept of crowdsourcing is now widely used for general AI dataset annotation (e.g., drawing a bounding box of a dog from the background), but paleontological studies often require more specific knowledge that the public is unlikely to have. Keçeli et al. (2017) manually segmented images of radiolarians and used AlexNet (Krizhevsky et al. 2012) to extract patterns from these manually processed features (e.g., eccentricity and circularity). They then used SVM, k-NN, Adaboost, and Random Forest to classify radiolarian species. The results showed that SVM generally performed the best among all classification methods. However, such a complicated workflow somewhat counteracts the idea of automation by AI. Since SVM had been applied to radiolarian classification previously by Apostol et al. (2016) and achieved equivalent performance, it may be unnecessary to apply CNN to extract data from manually processed features for subsequent classification.

Fenton et al. (2018) studied the consistency and accuracy of modern planktonic foraminifera identification. Experienced participants have higher accuracy than less experienced scientists, but surprisingly, participants can make good assessments about their own identifications, and the strongest predictors of accuracy are their knowledge and their confidence. Although working on modern taxa, the conclusion indicated the complexity of microorganism (and microfossil) identification. There is no doubt that machines can work much faster than any human, but how to evaluate the performance and the quality of training dataset should be more carefully considered.

Xu YX. et al. (2020) applied combinational machine learning including scale-invariant feature transform (SIFT), K-means, and SVM to automatically recognize microfossils from images, and compared the performances between these traditional machine learning methods and classical CNN models. The authors suggested that due to the existence of abiotic rock images, traditional machine learning methods were overwhelmingly better than CNNs, suggesting that CNNs may not be universally powerful especially when the tasks can be clearly delineated. As CNNs are usually very large models with millions to billions of parameters that need to be tuned during training, small datasets cannot fulfill the request to optimize all parameters.

#### 2.1.4 Other Deep Learning Methods and Dataset Construction

While model designs are undoubtedly important for AI studies, training data are also crucial for performance. Since paleontology suffers from data deficiency, most paleontological AI studies surveyed above were trained on only hundreds to thousands of samples, many of which were far from balanced (i.e., images are unevenly distributed among classes). Besides building larger datasets, another major strategy can be adopted to address this problem, *transfer learning*. Transfer learning uses a pre-trained model or part of it on new tasks. Depending on the relevance between the source and target domains, it saves training cost and takes advantage of relevant information from pre-training datasets.

Zhong et al. (2017) summarized previous work on foraminifera classification and proposed an automated pipeline for identification and discussed the limitations from computer vision in specimen identification including limited sample sizes and fixed observation angles. They used pre-trained classic networks (ResNet-50, VGG-16, and Inception V3) to save training costs in paleontological AI studies for the first time, indicating the versatility of neural networks and large datasets. Their dataset comprised 1437 images from 6 planktonic foraminifera species and was also studied by Ge et al. (2017) and Mitra et al. (2019). Keçeli et al. (2018) used *de novo* trained and pre-trained CNNs to classify radiolarians. Interestingly, the pre-trained VGG-16 models performed much better than the *de novo* trained CNN, but it is uncertain whether this was caused by neural network depth or pre-training. Mitra et al. (2019) trained CNNs to classify six foraminifera species and claimed that experts can achieve precision similar to a machine, but with lower accuracy. As Zhong et al. (2017) and Mitra et al. (2019) suggested, these AI models only performed a role as “proofs of concept” that CNNs can be used in paleontological studies, but are far from practical demonstrations of practical usefulness.

The *Endless Forams* dataset was created by Hsiang et al. (2019), including 34,640 planktonic foraminifera images covering 35 extant species. They showed that transfer learning based on classical datasets such as the ImageNet (Deng et al. 2009, including 14,197,122 images indexed by 21,841 synsets) performed well on the identification of foraminifera, achieving an accuracy of 87.4% using VGG-16. On the basis of the *Endless Forams* dataset and two other even larger datasets of benthic and planktic foraminifera from Holocene sediment cores (more than 600,000 images in total), Marchant et al. (2020) showed that larger training datasets and deeper neural networks can further facilitate foraminifera classification.

Karaderi et al. (2021) applied *deep metric learning* for the first time to the *Endless Forams* dataset to visualize the planktic foraminifer morpho-space structure. Deep metric learning is the learning of distance function between objects in high-dimensional metric space, and distance indicates the similarity in category or characteristic. In this case, the metric is the morphological distance between each pair of specimens in the *Endless Forams* dataset. The t-SNE visualization shows almost identical morphospace distributions between training and testing datasets, suggesting that larger datasets are more balanced and can result in better performances. By comparing with other published benchmarks on the same dataset, Karaderi et al. (2021) found that the deep metric learning exceeded other methods in identifying planktic foraminifera species, reaching an accuracy of 92%. However, the validity of metric learning and its natural deficiencies should be more carefully evaluated in comparison to the performances of different learning strategies (Musgrave et al. 2020). De Garidel-Thoron et al. (2020) presented microfossil sorter (MiSo) system that can automatically pick microfossils from other coarse sedimentary fractions and process up to ∼8,000 samples/day. Richmond et al. (2022) developed a system for foraminifera manipulation, sorting, imaging, and classification called Forabot. They used the dataset from Hsiang et al. (2019) and eventually the system can process approximately 27 specimen per hour, so it is more like a prototype than a practical tool. But the combination of hardware and software opened the possibility of fully automated data collection and analysis.

Liu X. & Song (2020) collected a large thin section image dataset of 30815 images from 18 fossil groups (mostly invertebrates) and four minerals or sedimentary structures. With pre-trained models on ImageNet, their results show high accuracies of >90%, and the more balanced dataset prevented overfitting in biotic structures. Following this, Liu X. et al. (2023) presented the Fossil Image Dataset (FID) with 415,339 images from 50 fossil clades (including various invertebrates, vertebrates, plants, microfossils, and trace fossils), and an online model (www.ai-fossil.com) is also available for fossil image identification. They showed that certain clades were more difficult to identify than others and fossil fragments hugely influence the rate of correct identification. Models pre-trained on ImageNet performed well regardless of the huge differences between ImageNet and fossil datasets. Moreover, activation mapping (the visualization showing image parts corresponding to the activation of the neural network) on fossil images may provide previously overlooked information about taxonomy and character evolution.

Pollen and other palynomorphs have commonly been explored using AI methods as specimens are abundant, similar to microfossil and insect studies. However, most palynological AI studies have focused on extant taxa, as reviewed by Treloar et al. (2004), Li P. et al. (2004), Zhang Y. et al. (2004), Holt et al. (2011), and Daood (2018). KBS, traditional machine learning methods, CNN, and transfer learning (Geus et al. 2019) have been applied in pollen image localization, recognition, identification, and classification. There are only a few AI studies on fossil pollen taxa and coverage has been restricted. Kong S. et al. (2016) proposed an unsupervised learning method to select representative feature patches from pollen images, then used these image patches as the dictionary as the basis of a sparse coding model. They performed SVM to classify pollen images into three selected species and reached an accuracy of 86.13%. Bourel et al. (2020) introduced CNN and decision trees in pollen recognition and claimed to be able to identify both fossil and modern pollens to the genus level, sometimes even species level. This study focused on three families (Amaranthaceae, Poaceae, and Cyperaceae) including 1698 pollen grains, in which 223 are attributed to the damage dataset and 97 are fossils. Although the integration of multi-CNNs and comparably large training datasets allowed successful identification of damaged pollen grains, the sampling strategy from only three families, a fairly short geological timespan, and geographically restricted localities in east Africa raise doubts about the generalizability of such a study. Besides taxonomic classification, Adaïmé et al. (2023), used various CNNs to extract pollen features, which are then passed to a multi-layer perceptron to predict phylogenetic distance from given taxa, last Bayesian inference was used to reconstruct the phylogeny. Adaïmé et al. (2023) limited their scope with the *Podocarpus* genus, but showed that deep learning can facilitate phylogenetic studies in character construction to uncover previously hidden evolutionary history. It may also provide insight for basic phylogenetic questions such as what a character is.

### 2.2 Macrofossil Classification

Macrofossils are large enough for direct observation by the naked eye, and are thus much fewer in number compared to microfossils. Their morphology is also more likely to be affected during preservation. Incompleteness and deformation of macrofossil specimens make it more challenging to construct a proper training dataset for automated classification. The classification of macrofossils often requires identification of diagnostic features, and they need larger data volumes to accommodate their morphological features. For example, a dinosaur skeleton 3D scan is much larger than a foraminifera 2D image. All these challenges have resulted in only a few examples of macrofossil classification studies, but there have been attempts to explore the potential of AI.

Based on a dataset from Huang & Shi (2022), including 16 fusulinid genera with 150 images of each, Hou C. et al. (2023) proposed a triple-base model using differently augmented images in the original, gray, and skeleton view (OGS) to improve identification performance. The OGS triple-base strategy showed generally better performance in almost all CNN architectures tested than using only one or two image types, and activation mapping indicates different hot-zones in different image types, indicating the complexity of characteristic features under different scenarios.

Insects occupy a large portion of biodiversity in both the modern and ancient biosphere, and the huge numbers of species and their widespread impacts in terrestrial ecosystems, as well the shortage of trained entomologists, have led to strong calls to identify and classify insects. Most entomological AI studies focused on extant species, of which Martineau et al. (2017) reviewed 44 studies for automated image acquisition, feature extraction, classification, and datasets. Many adopted traditional machine learning methods in those tasks while a few have tested the performance of neural networks. De Cesaro Júnioro & Rieder (2020) surveyed automated identification of extant insects with a primary focus on machine learning methods. Among the 33 studies examined, 63% used CNNs based methods in either recognition or classification tasks, while 29% used handcrafted features in the same tasks. Few studies introduced more complicated methods than CNNs, such as the attention mechanism. Contrary to foraminifera AI studies, large scale datasets have been better presented in extant insect studies. The largest dataset was proposed by Liu L. et al. (2019), the *PestNET* (https://www.pestnet.org/), which incorporated more than 80,000 images with over 580,000 pests classified into 16 classes (not necessarily species). *PestNET* not only utilized traditional CNNs for classification, but also introduced the attention mechanism in feature extraction and reached mean average precision of 75.46% in multi-class detection. There have been several other surveys and perspectives about AI applications in entomology (Valan et al. 2019; Høye et al. 2021, Kasinathan et al. 2021, Amarathunga et al. 2021). Since most insect-related machine learning studies worked with extant species, current AI-based entomological studies are probably not limited by the sparsity of data, but by the lack of well annotated datasets, incongruence in taxonomy, and currently overlooked links with molecular methods, such as DNA barcoding, that already provide plenty of successful AI applications.

Ho M. et al. (2023) combined prior knowledge hierarchical category relationships and CNN to classify organisms from petrographic images. They constructed TaxonNet based on Branch CNN that combined different branches with various classification categories to make predictions in a hierarchical manner. The results showed that the fine-to coarse Reverse TaxonNet can achieve the best accuracy of 90–95%, indicating that paleontological AI studies can be better facilitated by specially designed models according to the data.

Lallensack et al. (2022) discriminated ornithischian and theropod footprints using VGG-16 and a dataset with more than 1000 footprint outline silhouettes. The models performed better than five human experts on tridactyl dinosaur tracks within the scope of testing dataset (36 tracks), but sampling bias and information loss from 3D to 2D was a major problem. Wills (2023) compiled a theropod dinosaur tooth dataset including 1702 teeth and applied several machine learning methods to infer the morphotype of isolated teeth from Middle Jurassic localities. The results indicated that even isolated teeth bear enough information for classification to the level of family (e.g., Therizinosauroidea and Troodontidae), which can be assessed by machine learning methods.

Conceição et al. (2023) developed PaleoWood, a machine learning based classifier for Paleozoic gymnosperm woods. They sampled 62 genera of Paleozoic gymnosperm woods and 16 morphological characters for training and validation, and their models were based on logistic regression (LR), linear discriminant analysis (LDA), and k-NN. Convergence played a crucial role in influencing the classification results as the model cannot distinguish it from homology. Such a classification system is more like a re-run of KBS than more recent models.

### 2.3 Segmentation

Segmentation is a classic AI task that converts an image into partitions that represent the positions and shapes of certain objects (such as humans, cars, trees, etc.). Traditional segmentation techniques used thresholding to distinguish pixels based on their color values, and more complicated manual tools such as region growing were developed to work with more complex scenarios. However, methods based on linear interpolation are not capable of segmenting even relatively simple images. Another major pathway towards automated segmentation is edge detection, which is usually based on human-designed operators, such as the first-order Canny operator (Canny 1986) and the second order Laplacian of Gaussian (LoG) operator (Basu 2002). Because most of these operators have barely been applied in paleontological studies, we do not provide a more comprehensive introduction to these algorithms here. To our knowledge the only paleontological example using an edge detection algorithm at this time is FossilMorph, a program for fossil image statistical analysis and classification developed by Zheng et al. (2022). It made various measurements by edge detection operator processing on fossil images and used k-means clustering to categorize fossils into spheroidal, broken, and filamentous types. The popular CNN model was not used in this study and the classification was not based on a taxonomy framework, so that it may avoid bias from model design or prior knowledge.

The applications of various imaging techniques in paleontology have resulted in rapid growth in the availability of such data, covering both micro and macrofossils, and their modalities include optical micrographs, electron micrographs, and CT-scan slices. Traditional paleontological imaging segmentation is largely manually processed, thus requires tremendous efforts especially with high-resolution imaging techniques. While AI-based models have been widely applied in medical tasks covering almost all kinds of data modalities and tissues (Litjens et al. 2017, Shen D. et al. 2017), segmentation models were only developed recently for both 2D and 3D fossil images (Ge et al. 2017, Qin Z. et al. 2022, Yu C. et al. 2022).

Ge et al. (2017) used the segmentation of foraminifera images by random forest and CNNs to differentiate apertures and different chambers in isolated foraminifera individuals. Since the morphology of apertures and chamber arrangements are keys to foraminifera species identification, this study may facilitate data labelling and species identification.

Hou Y. et al. (2020 & 2021) published ADMorph (Archives of Digital Morphology, http://www.admorph.ivpp.ac.cn), an open dataset of vertebrate fossils for deep learning studies and tested the segmentation performance of various classical DNNs such as U-net, PointNet, and VoxNet on that dataset. These two studies show that paleontological data can be prepared and processed in the workflow resembling other kinds of images and deep learning can significantly save processing time in imagery tasks such as segmentation.

Yu C. et al., (2022) reported the segmentation of dinosaur fossil CT scans by deep learning^3^. This study constructed a dataset comprising more than 10,000 labelled CT slices derived from three embryonic *Protoceratops* skulls discovered in Mongolia. Such fossils, at early developmental stages, are poorly ossified and difficult to differentiate from rock matrices. Skip connections and Atrous Spatial Pyramid Pooling (ASPP) were used to capture features at different scales. Although this model performed well in the given datasets, it is poor in general when segmenting fossils from other organisms and sedimentary environments.

Qin Z. et al. (2022) studied the distribution of osteons in a group of highly specialized theropod dinosaurs, Alvarezsauria. Histology studies the structure and morphology of tissues, mostly bones in vertebrate paleontology, to provide information about the growth, development, physiology, and even reproduction of animals. Alvarezsaurian dinosaurs underwent rapid size miniaturization during their evolution (Qin Z. et al. 2021), but the pattern of such change in body size remains unknown. There are usually two pathways toward miniaturization: truncated development and compressed development through the early termination and shortening of development, respectively. Qin Z. et al. (2022) sampled six alvarezsaurian taxa across the evolutionary history of this group, covering large, medium, and small body sizes. By applying a dual-resolution segmentation strategy, the DNNs used were able to efficiently localize the central vascular canal of primary and secondary osteons, then expanded to cover the entire osteon structure. By comparing the number and area of primary and secondary osteons in different taxa, they suggest that the miniaturization in Alvarezsauria was most likely to have been caused by truncated development.

### 2.4 Prediction

Prediction is also a major task in AI applications, namely, to estimate the probability or continuous value of a certain output according to the intrinsic laws learned from training datasets. A typical example of AI prediction in earth sciences is weather forecasting. By integrating historic patterns and data from surrounding regions, AI can make short-term prediction on future local weather with reasonable accuracy (Reichstein et al. 2019; Bi et al. 2023; Zhang YC. et al. 2023). Although a substantial number of paleontological studies focus on prediction of certain aspects of extinct organisms (e.g., their behavior, soft tissue morphology, and ecology), AI has only been applied in a limited way to such studies.

While AI application to fossil insect classification is limited (section 2.2), there have been a few paleo-ecological studies using AI to predict ancient mimicry behaviors. Fan L. et al. (2021) studied plant mimesis in both extant and extinct insects, which showed similar results consistent with the deep origin of biological mimesis. Based on the same neural network, Xu C. et al. (2022) further analyzed mimicry and insect camouflage from mid-Cretaceous Kachin ambers. In these two studies, the Siamese Network was firstly pre-trained on the Totally-Looks-Like dataset (TLL, Rosenfeld et al. 2018), which comprises 6016 image pairs that look similar to the naked eye but may come from totally unrelated objects, then the model was fine-tuned by specifically constructed mimic insect-plant pair datasets. Dissimilarities are measured between plant-insect pairs to quantify mimesis behavior. However, both studies were confined by fine-tuning of the datasets as they could not carry out exhaustive sampling over all plant-insect mimesis behaviors. Nicholson et al. (2015) suggested that there had been a fast expansion in our knowledge of extinct insect diversity and taxonomy since 1994. Established fossil insect databases, although not as comprehensive as extant insect databases, are ample resources for data-driven entomological studies, particularly AI-based studies.

Anemone et al. (2011) made attempts to use AI for fossil exploration prediction. A four-layered ANN was trained to explore the connection between landscapes and fossil preservation, and the results from Bison Basin, Wyoming, USA seemed to be encouraging. Martín-Perea et al. (2020) showed AI-based prediction of fossiliferous levels in both archaeological and paleontological sites, with testing results from two Late Miocene paleontological sites in Madrid, Spain. Multiple methods including SVM and RF were used to quantify the spatial distribution of fossils and these provided information about the faunal assemblage and directions for excavation.

We cannot discuss every paleontological AI study in detail. Fig. 3 illustrates major progress in the development of paleontological AI and mainstream AI. An introductory list is presented in Table 1 with publication dates, study organisms, methods, and tasks. In the following two sections, we present several paleontological AI studies that do not belong to the previous four categories but may provide insight into general AI applications in paleontology.

**Figure 3.**
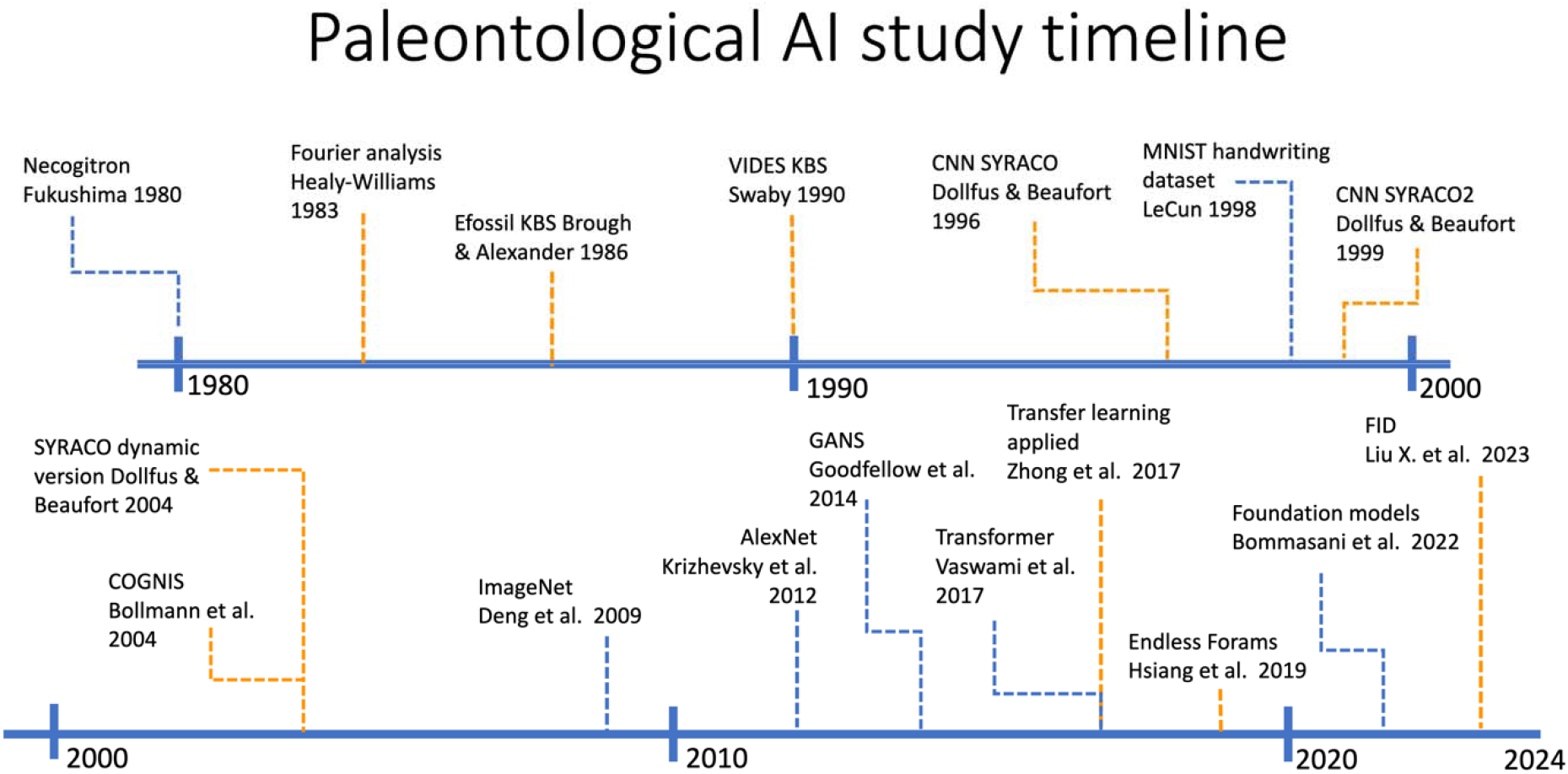
Major progress in paleontological AI study.

**Table 1.**
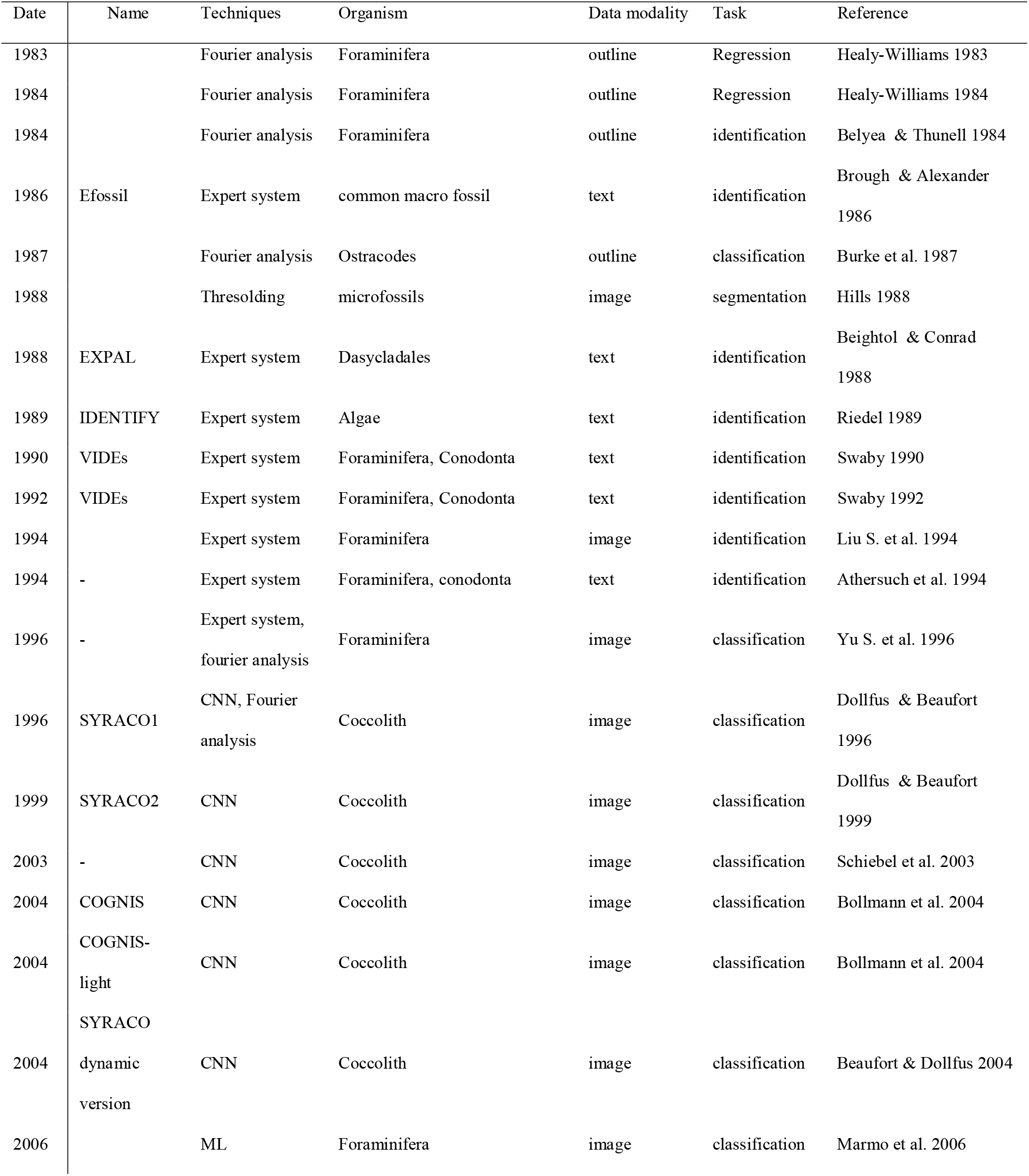
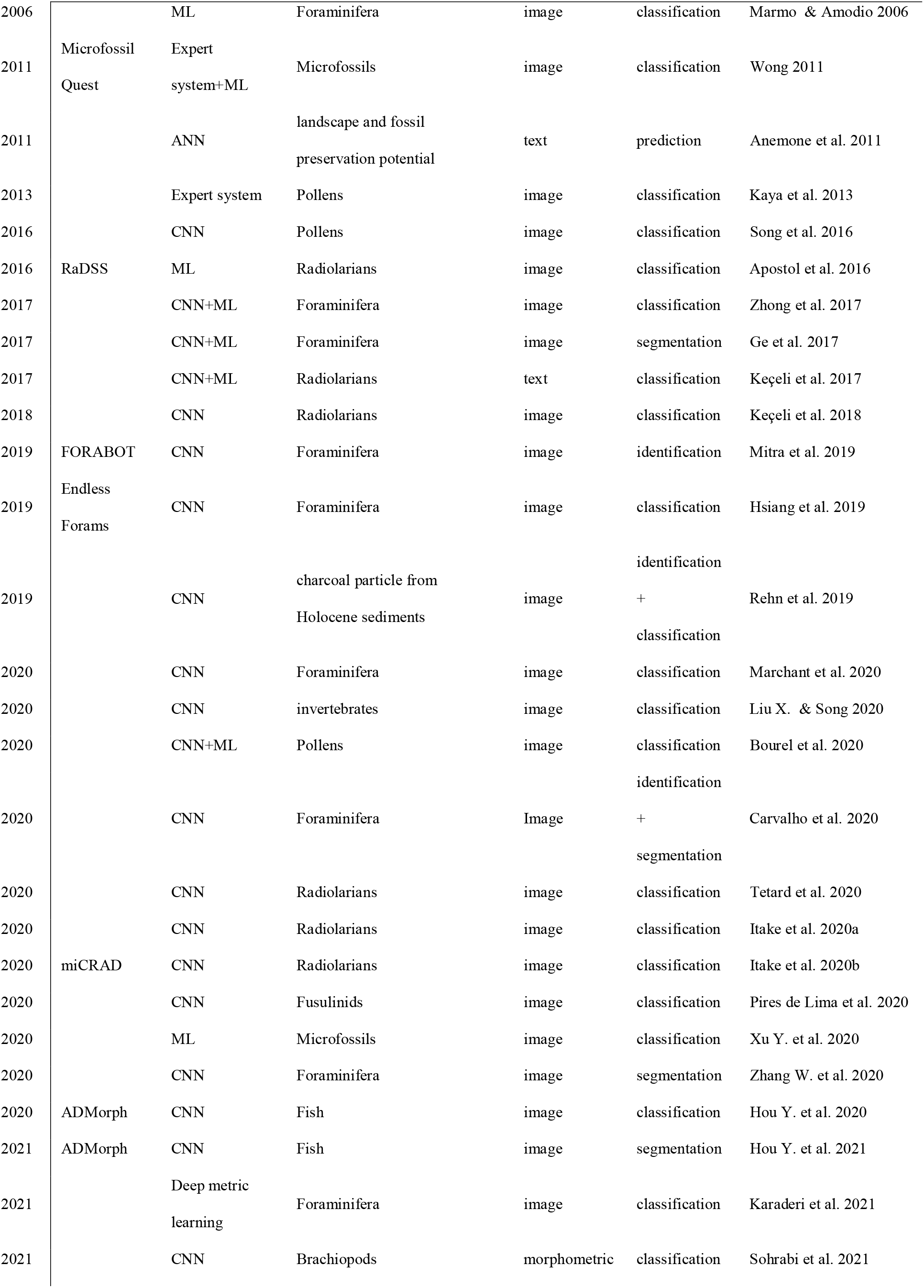
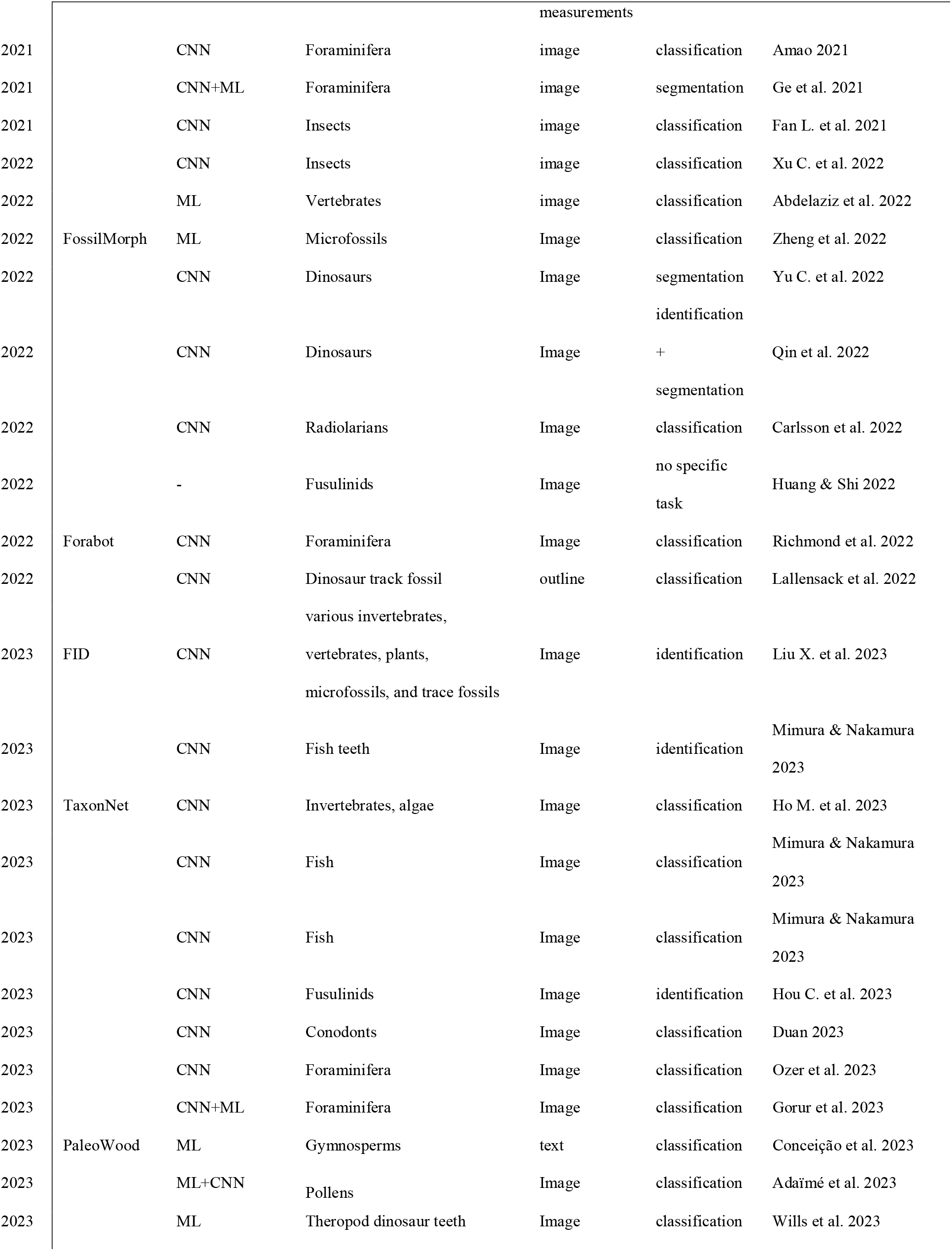
A list of paleontological AI studies in chronological order.

### 2.5 Summary

Here we reviewed paleontological AI studies primarily by their tasks (classification, segmentation, and prediction), Among the studies surveyed here, roughly 1/3 focused on foraminifera, 1/3 on other microfossils, and the rest on other organisms including plants, insects, dinosaurs, etc. (Fig. 2B). A possible explanation for such preference is data availability. Microfossils (including foraminifera) can often be completely preserved in marine sediments and suffer less deformation compared to macrofossils. And many microfossils have been key indicators for paleoclimate, paleogeography, stratigraphy, and many other research fields. The need to process large quantities of specimens quickly has driven the development and deployment of microfossil AI applications. Most practical AI applications (SYRACO and COGNIS) and open datasets (*Endless Forams* and FID) have focused on microfossils to date. Indeed, all AI methods mentioned above have been applied in microfossil classification; it is thus a representative example of the unfolding development of paleontological AI studies. Studies on macrofossils are notably fewer, with most working with CT scans if individual structure/elements and histological thin sections, which are not strongly relevant to gross fossil morphology and taxonomic identification. Liu X. et al. (2023) built one of the largest paleontological AI datasets, FID, covering various macrofossil clades. However, their results showed that AI performance on microfossils is generally better than on macrofossils.

While microfossils have been the most popular organisms in paleontological AI studies since the 1980s, studies on other small-sized fossils like insects and plant pollen are rare despite their large quantities of available fossils. Various reviews have summarized AI applications on extant insect and plants (Treloar et al. 2004, Li P. et al. 2004, Zhang Y. et al. 2004, Holt et al. 2011, Martineau et al. 2017, Daood 2018, and De Cesaro Júnioro & Rieder 2020), showing that most studies worked with extant species probably due to better annotated datasets and their practical significance. There is also interoperability between AI identification and classification for extant and extinct taxa. Rani et al. (2022) reviewed advances in automatic (extant) microorganism recognition, among 100 studies from 1995 to 2021 and, although only a single paleontological study (Mitra et al. 2019) was mentioned, they showed similar trends in the development of methods and data scales.

## 3. Results

### 3.1 General Trend

While the earliest paleontological AI study can be traced back to the early 1980s, and relevant discussions appeared even earlier (Sneath 1979), there was <1 published study per year on average before 2000. The number of studies and data scale only significantly increased recently (Fig. 4). There have been remarkable changes in the methods used, while the studied organisms, tasks, and data modalities have remained stable through the last four decades.

**Figure 4.**
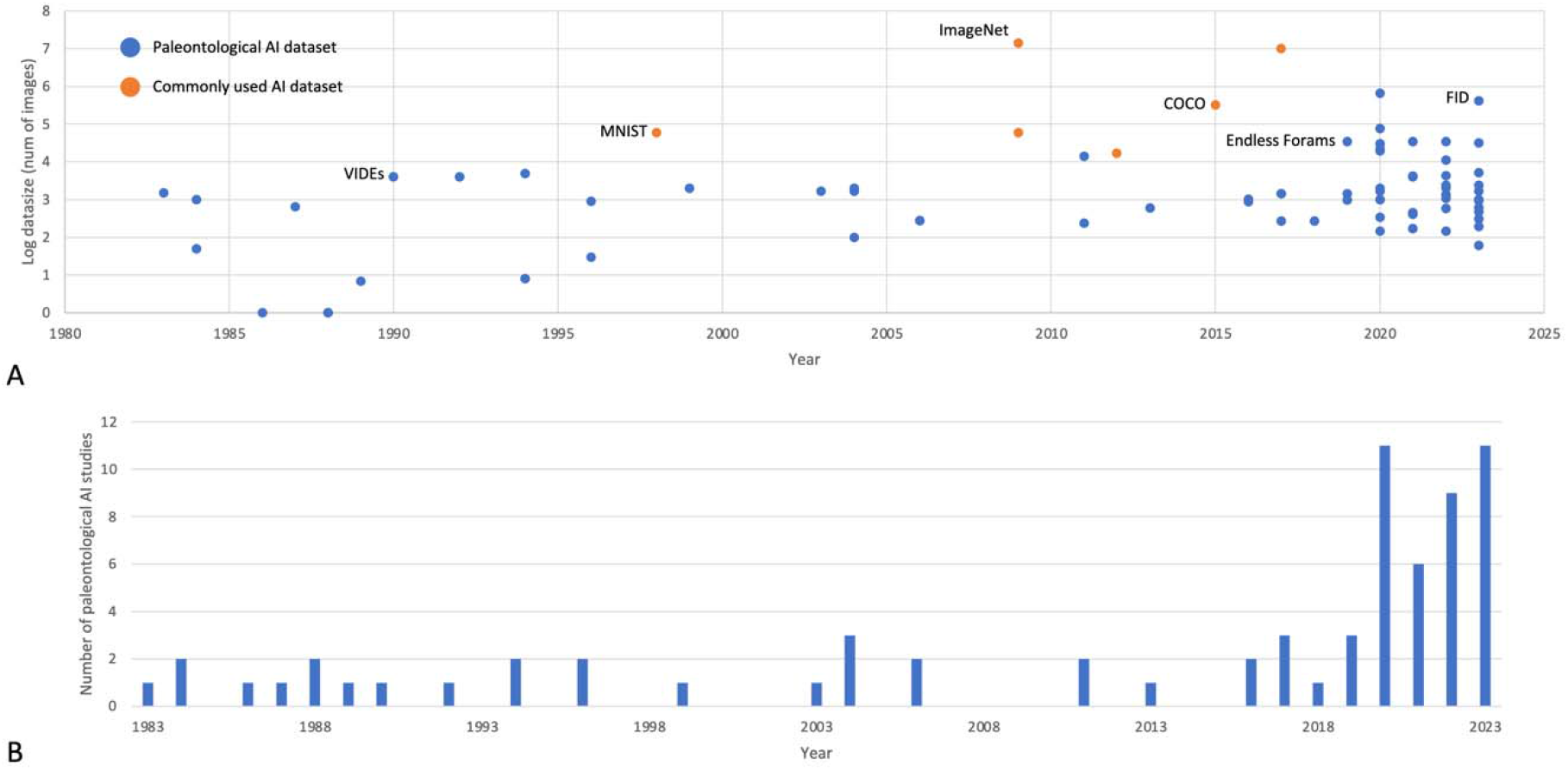
A. Data size (log10) of paleontological and commonly used AI datasets from 1983 to present. B. Number of paleontological studies from 1983 to present.

### 3.2 Methods

During the 1980s to mid-1990s, the first generation of fossil AI was based on KBS, resembling the evolution of AI models at the same time or slightly earlier. But KBS, normally as a handcrafted rule-based system, has internal deficiencies in both system building and maintenance, and was rapidly replaced by CNN in general AI applications and more specifical fossil studies. CNN has proven to be effective in imaging and a variety of other tasks. Although the first attempt building CNN-based AI model was by Fukushima (1980), its first application in paleontological studies occurred much later (Dollfus & Beaufort 1996).

CNN has been the most common model in current paleontological AI studies. It is the sole model used in more than half of the studies surveyed here, even without counting those incorporating multiple models (Fig. 2B). With the maturation of opensource machine learning frameworks such as TensorFlow and PyTorch, building neural network models is becoming easier. And the fast advancements in GPU and availability of cloud computing services like Google Colab allow almost anyone to access unprecedented computational power. There are also plenty of ready-made models for training or fine-tuning. Most *de novo* designed CNN models in paleontological AI studies were shallow, seldom exceeding five layers. Meanwhile, AlexNet (Krizhevsky et al. 2012) had five convolutional layers and three fully connected layers, and Deep Residual Network can reach hundreds to thousands of layers in depth (He et al. 2016). There is a significant depth gap between proposed paleontological AI models and classic works. Part of this is that the amount of data needed to train very deep networks is not available in current paleontological studies. Hsiang et al. (2019) tested VGG-16, Inception V3, and DenseNet-121 on the Endless Forams datasets, which significantly outnumbered most paleontological datasets surveyed here, but the shallowest VGG-16 had the best performance. Many researchers have admitted that the paleontological AI models they developed were only “proof-of-concept”, which refers to both models and data coverage. At the moment, we have witnessed success from much more complicated models on complicated tasks, and there is now hardly any need to validate the feasibility of CNN or many other models in paleontological AI studies.

Transfer learning has also been widely used to overcome the computational cost at training stage. Many studies chose to use models pre-trained on large image datasets such as ImageNet (Deng et al. 2009, Zhong et al. 2017, Keçeli et al. 2018; Mitra et al. 2019, Hsiang et al. 2019, Marchant et al. 2020). Classic networks including VGG-16, ResNet50, and U-net are often preferred as the models or at least the backbones of larger models. There have been very few specifically designed models for paleontological AI studies (e.g., Marchant et al. 2020 created a custom CNN, Base-Cyclic, that adapts to image size; Qin et al. 2022 designed a dual-resolution network to distinguish primary and secondary osteons in histological thin sections), and in any case these may not be necessary.

While CNN is the most favorable model in paleontological AI studies, traditional machine learning methods including SVM, k-NN, k-means, RF are still actively conducted. Xu YX. et al. (2020) criticized the overuse of neural networks in fossil image recognition. In spite of its advantages, CNN-based models usually need to be trained on large and balanced datasets, and the results are often fragile in interpretability. Paleontological data are intrinsically sparse and fragmentary; as a result, CNN may be outperformed by other machine learning or even more concise methods under many scenarios.

### 3.3 Tasks and Data Modalities

About two-thirds of paleontological AI tasks are multi-class classification as it plays a fundamental role in paleontology (Fig. 2B). The established taxonomy offers a solid basis for training supervised learning models, thus in principle we can construct model that is capable of accommodating all taxa of interest. However, the problem is that the amount of training data that’s necessary to train such a model is largely unfeasible to collect. Other typical tasks include (image) segmentation and prediction, while they are both classic AI applications, the number surveyed here is significant fewer than classification task. Other common tasks in mainstream AI studies, such as text processing, image inpainting, and feature engineering, remain untouched in the paleontology field.

Around 80% of studies surveyed here used image datasets, and several had outlines or silhouettes that are directly derived from images (Fig. 2B). This is not surprising since images are the most direct and inexpensive method to record fossil morphology, especially for microfossils. However, the morphology of macrofossils cannot be easily captured by such low-resolution 2D images. A detailed 3D image that is enough for morphology description, for example a high-resolution CT scan, is too large for current AI model training. There are also too few of them available in general to adequately train models. In practice, high-resolution macrofossil images are often used for illustration, measurement, morphology description reference, and geometric morphometric data collection, but not directly in analysis. Given the current computational capabilities, the large 3D and high-resolution 2D images need to be compressed for subsequent studies. There have not been paleontological AI studies working with those compressed data modalities from original images, and the compression process itself may be a target for AI learning.

### 3.4 Data Scale

Large amounts of data are necessary for AI model training. Early paleontological AI studies were usually based on KBS, and datasets comprising up to ∼10^3^ images were constructed in 1980s (Swaby 1990, Fig. 4). Later CNN-based models were often trained on hundreds to thousands of images. The construction of large-scale open datasets, *Endless Forams* for example, was a notable advance. Several fossil image datasets have reached ∼10^4^ to 10^5^ images, but many recent studies continue to use small datasets with fewer than 10^3^ images (Fig. 4). Although the incremental progress and proof-of-concept studies are limited in their practical applicability, without innovate methods, those small-sample studies may claim to have equivalent or even better performance than human, but are far from practical usage or scientific significance. The recent boom in number of paleontological AI studies is more likely the result from a continuously lowering bar in training and deploying AI models than real progress, at least from the perspective of data scale.

The scales of current foraminifera datasets have reached or even exceeded classical deep learning datasets such as the MINST handwritten digit dataset (http://yann.lecun.com/exdb/mnist/, 50,000 training and 10,000 testing images in 28 × 28 pixels) created in 1998, but still are 2∼3 orders smaller than other commonly used baseline datasets (ImageNet has 14 million images in >20,000 categories). There is roughly a 20-year lag between mainstream and paleontological AI studies in the term of data scale.

Most datasets surveyed here were manually collected and annotated, but there are also exceptions by using web crawling and published literatures (Liu X. et al. 2023) for collection and crowdsourcing for annotation (Wong 2011), which are both common practice in preparing large-scale AI training datasets. However, paleontology is a specialist field that needs expert knowledge and long-term practice in order to generate training sets, making automated and crowdsourced dataset creation a more difficult prospect. There have not been unsupervised learning examples in relevant studies, and relying on automatically crawled data or crowdsourcing can cause unnecessary bias during model training.

## 4. Perspectives

In this section, we discuss the various aspects of paleontological studies that can have benefited from established AI achievements in foreseen future, and how recent progress including Transformer, one/few-shot learning, auto-content generation, and Large Language Models will interact with paleontology.

### 4.1 Automated Workflow

Automated species classification has always been a focus in paleontological AI studies, such idea was repetitively proposed by Gatson & O’Neill (2004), MacLeod et al. (2010) and many others, and now there is prototype system developed based on large datasets and deep learning techniques (Liu X. et al. 2023). From KBS to CNN, most (if not all) paleontological AI studies surveyed here are within the range of supervised learning, meaning that there needs to be a reliable labelled dataset for model training, with clear taxonomy for fossil classification.

Taxonomy and phylogeny of extinct organisms is fundamentally based on their morphology, more specifically synapomorphies in the context of cladistics. However, the definition of most morphological characters is fundamentally qualitative and subjective, as is the interpretation of homologies – not at the analysis level, but at the character definition and coding level. There is good reasoning and evidence for many characters, but that doesn’t change the fact that it’s a qualitative/subjective determination by human, and different researchers can and do disagree about different hypotheses of characters/homologies. Linear classifiers on top of hand-engineered features are only able to make simple partitions of the output space, indicating more complex models are obligatory for automated species classification.

To avoid personal bias, artificial (morphological) characters/features will probably no longer be the foundation of reconstruction fossil species taxonomy or phylogeny. Semi-supervised or even unsupervised learning can facilitate the discovery of synapomorphies and diagnostic features. The application of transfer learning since 2017 has indicated the feasibility of automated character construction as models pre-trained on common objects showed appreciable performances on fossil datasets. Activation maps, which show the regions of attention that the machine uses to make its determinations, have been applied to fossil image classification (Hou C. et al 2023; Liu X. et al 2023) and suggest that machines are capable of discovering previously hidden patterns in taxonomy.

Even though automated classification sounds unrealistic at this moment for paleontologists, AI can certainly help to automate current paleontological workflow. The implementation of AI into GM has shown exciting potential for high throughput phenotypic research. Studies have used machine learning algorithms for classifying existing shape data into clusters (e.g., Soda et al. 2017; Courtenay et al. 2019; Arriaza et al. 2023). While classification of data is a valuable application of AI, the greatest gains will likely come from using AI to automate the collection of quantitative phenotypic data. Collection of landmark-based geometric morphometric data is largely done manually and often a rate limiting step in current phenotypic studies. (Semi-)automated landmarking tools exist, including TINA Geometric Morphometrics Tool (Bromiley et al. 2014), auto3DGM (Boyer et al. 2015), ALPACA (Porto et al. 2021) implemented in the open software SlicerMorph (Rolfe et al. 2021), and others (e.g., Aneja et al. 2015). Alternatively, users can employ landmark-free methods that permit visualization and analysis of shape variation based on 3-D mesh models, such as Generalized Procrustes Surface Analysis (Pomidor et al. 2016), spherical harmonics (Shen L. et al. 2008; Dalmasso et al. 2022), non-rigid surface registration (Snyders et al. 2014; Claes et al. 2018), and Morphological Variation Quantifier (morphVQ; Thomas et al. 2023). Although these techniques empower users to rapidly collect rich phenotypic data, AI has the potential to further automate and enhance these efforts.

To date, ML applications for phenotypic data have largely proceeded on clinical data and human faces. For rapid phenotyping and analysis of diverse biological systems, both supervised and unsupervised machine learning approaches could become indispensable. Currently, ML-morph (Porto & Voje 2020) and another tool (Vandaele et al. 2018) provide a supervised approach for automatically placing landmarks on 2-D images. These studies examined the performance of these methods on sets of different biological structures, ranging from *Drosophila* wings to entire bodies of fish, exhibiting comparative variation to manually landmarked datasets. Because these are supervised ML approaches, training datasets are required that consists of corresponding image and landmark data for dozens, if not hundreds, of specimens. Wöber and colleagues (2022) used CNN on 2-D images of fishes to allow for major anatomical variation to be identified without *a priori* decisions on morphological features of interest. The authors report that the morphological features picked up by CNN is similar to what the manually collected landmark data show as distinctive areas between fish populations and that it was able to identify distinct groups that match population clusters based on genetic data than landmark data (Wöber et al. 2022). Existing ML tools show much promise for automated landmarking with high accuracy and precision under supervised algorithm, whereas unsupervised algorithms could lead to new discoveries about phenotypic differences and transformations. Extensions of these methods to 3- D image data (e.g., µCT data) will be paramount for our efforts to characterize phenotypes of larger organisms (e.g., vertebrates) across the tree of life.

Despite enormous potential of ML approach for GM, ongoing utilization and future advances in these techniques require important considerations. First, image acquisition standardization will be critical for both supervised and unsupervised ML methods because the accuracy of phenotypic characterization will be strongly influenced by image properties and quality. One way to reduce these effects is to expand the training dataset to include images in variable settings and orientations, although this strategy come with obvious costs. Extending the implementation of ML tools to 3-D surface data may allow to readily consider differences due to orientation. Secondly, as with any landmark data, consideration of landmark scheme that suits the biological question of interest is crucial. ML methods, particularly unsupervised approaches, involve greater layer of model assumptions and may select morphological features that may not be biologically significant to human (e.g., lack of evolutionary homology). What one gains in speed and automation may lead to loss in interpretability. Thirdly, investigators should ensure replicability of ML methods by reporting on any parameters used for the AI procedure and making any training datasets openly available. Next, these AI tools should allow for landmark positions to be checked and adjusted manually, ideally within the same program (e.g., Bromiley et al. 2014; Vandaele et al. 2018). This allows an investigator to easily assess the quality of the landmark placement and revise as needed. Finally, advancements in ML techniques, as well as non-AI automated landmark approaches, that allow for accurate phenotyping of broad, comparative sampling or developmental series should be made. Thus far, these methods have been applied to fairly restricted datasets, mostly within a genus.

### 4.2 Models and Datasets

There is roughly a 20-year latency in paleontological dataset scale comparing with mainstream AI studies, while the latency is shortened to about 10 years with respect to model usage given that AlexNet was proposed in 2012 (Krizhevsky et al. 2012). Currently only basic CNNs and several primitive variants have been applied to fossil data. Nevertheless, even a simple CNN architecture can largely outperform many other methods. More importantly, it has been realized that, generally, increasing neural network depth can rapidly approximate extremely complicated functions that are challenging to be described using handcrafted features.

Deep learning technology is still developing rapidly. The progress of DL mainly lies on three aspects: base model, task head, and learning paradigm.

1. The base model acts as the backbone of a deep neural network, which determines the feature representation ability. The early AlexNet has a simple structure, i.e., only five convolutional layers (Krizhevsky et al. 2012). In later years, base models with more complex structure were proposed. ResNet expands the network depth up to 200 layers by introducing residual unit (He et al. 2016), which significantly improves the feature extraction ability and training convergence. Beyond convolutional neural networks, Vaswani et al. (2017) introduced Transformers, which use a self-attention mechanism that weights different parts of inputs based on their significance. The novel attention-mechanism-based Vision Transformer (ViT) and multi-layer perceptron based MLP-Mixer, which achieve promising performance when the pre-training dataset is large enough, have also been proposed (Dosovitskiy et al. 2020, Tolstikhin et al. 2021). Although the base model is general purpose, specific base models are often required in specific tasks. For example, the stacked Hourglass Network was proposed specifically for keypoint detection (Newell et al. 2016), and dynamic graph CNN was proposed specifically for point cloud encoding (Wang Y. et al. 2019). Given the uniqueness of paleontological data, the design of specific base model is of considerable importance.
2. The task head is used to process the features offered by the based model and then, infer the output. The design of task head is task-specific and determines the output data’s format. The classification task head is very simple, which can be realized by simply pooling the feature map and feed the resulting vector to a classifier. The task head design becomes more challenging when describing the object’s details using fine-grained outputs (Qin F. et al, 2022). In many cases, the DL model has multiple output branches to fulfill multi-task inference (Kim et al. 2022). Multi-task prediction and fine-grained outputs are also required in many paleontological tasks.
3. The learning paradigm refers to how the model is trained. The most widely used paradigm is the supervised learning on a large annotated dataset. The design of the training loss function significantly influences the training outcome (Wang Q. et al. 2020). Since data collection and annotation are expensive and time-consuming, which is especially the case in paleontology, large data driven supervised learning can hardly be feasible. Therefore, a new learning paradigm is urgently needed to overcome the data sparsity problem. Many researchers have been working on *few-shot learning*, aiming to enable the AI model to learn rapidly from only a few known examples, just as humans learn (Wang YQ. et al. 2020). Few-shot learning has been explored in many scenarios, such as CT medical image diagnosis and microscopic vision measurement (Chen et al. 2021 & Qin F. et al. 2023). Further, semi-supervised learning uses both labeled and unlabeled data for learning, which can also relieve the data sparsity problem and exploit huge amounts of raw data (Van Engelen et al. 2020). In the future, we can expect more novel, specially designed AI models and high efficiency but low-cost learning paradigms for paleontological data processing.

Data availability is another significant obstacle for future paleontological AI studies. Existing studies often come with a tremendous waste of data. A typical CT-scan of a dinosaur skull may result in a 3D model of several gigabytes, but after encoding it into a morphological character matrix, the overall morphology turns into tens to hundreds of bytes, whose compression rate is approximately 10^6^. Such drastic waste of data very likely results in the loss of relevant information. Moreover, the coding process is often subjective, being based on an understanding of anatomy and evolution. Several studies (e.g., Allmon et al., 2018) suggested that paleontology is going to embrace Big Data. However, Big Data should usually meet the requirements of the four Vs: volume, variety, velocity, and veracity. Paleontological data have deficiencies in at least the volume and collecting velocity terms. Among recent data-driven studies, only a few have worked with data in a non-handcrafted manner (e.g., Fan J. et al., 2020), while most of them, are still within the frame of traditional handcrafted workflows (though with larger datasets than studies before, e.g., character matrices containing thousands of characters). In the next decade or two, we may expect that many recently proposed methods in AI will be transplanted to paleontology and there will be larger, better labelled, better organized and maintained datasets for both model training and further studies. It may be the beginning of the transformation towards data-driven paleontological studies.

### 4.3 Multi-Modal Model

Currently paleontological AI models have only been developed for restricted tasks. However, realistic paleontological studies often gather various data modalities and associated metadata including fossil morphology, geological age, paleo-environments of localities, isotopic composition, small or even macro molecular remnants, etc. Fossil data collection is not limited to standard images and text, but rather encompasses images from various advanced imaging techniques, the occurrence information of fossil localities, morphological measurements, morphological character matrices, phylogenetic trees, and contents from published literatures, most of which remained untouched by AI models but will surely be a goldmine for data-driven studies.

Recently, promising progress has been made beyond applying AI to particular tasks, but to create unified models to integrate multi-modal data (e.g., text, images, sequences, etc.), to accomplish various tasks (Radford et al. 2021). Models such as Bootstrapping Language-Image Pre-training (BLIP, Li J. et al., 2022) and Once for All (OFA, Wang P. et al., 2022) show that multi-modal models can reach comparable performance with uni-modal models in not only classic machine learning datasets, but also unseen tasks and domains. The combination of image and text data is capable of automated generation of text based on certain images and identify objects with certain description from images, which is similar to a major task in traditional paleontology, namely the description of fossil morphology. Fossil description occupies a fundamental status in paleontological studies and traditionally requires long-term training in anatomy, systematics, physiology, and many other areas. The accomplishments achieved by these large pretrained models may pave the way towards the automated description of fossils in the future, which may lead to fundamental changes in the paradigm of paleontological studies since the birth of this field.

### 4.4 New Techniques

There have been many innovative and successful methods that are totally unfamiliar to paleontology, such as Generative Adversarial Networks (GANs, Goodfellow et al. 2014), diffusion models, and Large Models.

GANs usually have two neural networks, a generator and a discriminator, that compete with each other to produce outputs that are as close to “realistic” samples as possible, which means they can be trained in both supervised and unsupervised manner. One of major applications of GANs is content synthesis and manipulation that can possibly be used for missing paleontological data complementing, which is similar to image inpainting (Guillemot & Le Meur 2014; Elharrouss et al. 2020). GANs belong to a large family of models called generative models, to which diffusion models (which outperform GANs in image synthesis) also belong (Sohl-Dickstein et al. 2015, Ho J. et al. 2020). Diffusion models are trained to learn the structure of training datasets in the latent space, similar to the diffusion process in physics. They are then able reverse the process to conduct tasks such as, denoising images. The recently developed content generation systems DALL-E (https://openai.com/product/dall-e-2) and Stable Diffusion (https://stability.ai/stable-diffusion) have shown astonishing performance in text-to-image generation using diffusion models, and can potentially be used for more complicated and customized content such as videos.

Most (vertebrate) skeletal fossils are incomplete and soft tissues can hardly be preserved in fossils. As such, the original organisms have to be reconstructed based on the relevance between anatomical structures, environmental constraints, and morphology from closely related taxa, etc. Such reconstruction is necessarily influenced by subjectivity. Further, along with the reconstruction of fossil organism themselves, the reconstruction of phylogenetic relationships is often impeded by missing links in evolution resulting from the patchiness of the fossil record. AI based generative models including GANs and diffusion models provide the chance to better hypothesize even reconstruct what has been lost from fossil preservation on the basis of more comprehensive scope and reproducibility.

Since the proposal of the attention-based transformer by Vaswani et al. (2017), its parallelization has led to large pre-trained models such as BERT (Bidirectional Encoder Representations from Transformers, Devlin et al. 2018) and GPT iterations (Generative Pre-Training Transformer, Radford et al. 2018 & 2019, Brown et al. 2020; OpenAI 2023), also called Large Language Models (LLMs). LLMs often have more than one billion parameters, dwarfing complexity of current paleontological AI models. Many transformer-based models perform better than traditional CNNs in various tasks such as image classification, object detection, language translation, etc. Due to their unprecedented large sizes and power, LLMs, or Foundation Models, and their concomitant training costs and emergent properties, have hugely impacted AI development and applications (Bommasani et al. 2022). Although none of those models have yet been tested on paleontological tasks, their prospective impact on paleontology may be greater than all other AI techniques we discussed above.

## Authorship contribution statement

Conceptualization: CYY

Investigation: CYY, FBQ, AW, WQY, YL, ZCQ, YML, HBW, QGJZ, AYH

Project administration: CYY, CM, ER, MJB, XX

Visualization: CYY

Writing original draft: CYY, FBQ, AW

Writing reviewing & editing: Everyone

## Data availability

All data used in this research are described in this article.

## Acknowledgments

We would like to thank Mark A. Norell and Jin Meng from the American Museum of Natural History for their valuable discussion at early stage of this project. This work is funded by the National Natural Science Foundation of China (Grant No. 42288201, 42272017); the Youth Innovation Promotion Association, CAS (2021068); Yunnan Revitalization Talent Support Program (202305AB350006); Swedish Research Council (Vetenskapsrådet) Starting Researcher Grant (ÄR-NT 2020-03515). This work is a contribution to the Deep-time Digital Earth (DDE) Big Science Program.

